# Density Physics-Informed Neural Network reveals sources of cell heterogeneity in signal transduction

**DOI:** 10.1101/2023.07.31.551393

**Authors:** Hyeontae Jo, Hyukpyo Hong, Hyung Ju Hwang, Won Chang, Jae Kyoung Kim

## Abstract

The transduction time between signal initiation and final response provides valuable information on the underlying signaling pathway, including its speed and precision. Furthermore, multimodality in transduction-time distribution informs that the response is regulated by multiple pathways with different transduction speeds. Here, we developed Density physics-informed neural network (Density-PINN) to infer the transduction-time distribution, challenging to measure, from measurable final stress response time traces. We applied Density-PINN to single-cell gene expression data from 16 promoters regulated by unknown pathways in response to antibiotic stresses. We found that promoters with slower signaling initiation and transduction exhibit larger cell-to-cell heterogeneity in response intensity. However, this heterogeneity was greatly reduced when the response was regulated by slow and fast pathways together. This suggests a strategy for identifying effective signaling pathways for consistent cellular responses to disease treatments. Density-PINN can also be applied to understand various time delay systems, including infectious diseases.

## Introduction

Cells respond to signals from their extracellular environment through complex intracellular signaling pathways. While reliable signaling is necessary for proper cell function, the timing and strength of the response to the same extracellular signal can vary significantly, even in genetically identical cell populations.^1-9^ This cell-to-cell heterogeneity leads to the emergence of abnormal cells, which can cause disease. Furthermore, heterogeneity can lead to incomplete killing of target cancer cells^10^ and the emergence of persister cells^11^, which are major obstacles in effectively treating cancer.

Previous studies have focused on indirect sources of cell-to-cell heterogeneity in signaling responses,^12-15^ such as cell cycle phase ^16^ and RNA polymerase level.^17^ On the other hand, direct sources, i.e., the signaling pathways themselves, have been investigated in a limited number of studies when comprehensive information about the pathways is known.^18-20^ For example, Granados et al. modified known signaling pathways leading to Hog1 expression and found that yeast shows low cell-to-cell heterogeneity in response to osmotic stress when the response is regulated by multiple pathways with both slow and fast signaling transduction speeds.^18^ Chepyala et al. developed a mathematical model for a known regulatory pathway in Caenorhabditis elegans and investigated the role of the pathways in controlling the heterogeneity of distal tip cell migration timing.^20^ However, comprehensive information about signaling pathways is rarely known except for these limited cases, rendering it challenging to identify the sources of the cell-to-cell heterogeneity within the signaling pathways themselves.

To overcome this lack of information about signaling pathways, one promising solution is to develop a model by replacing an unknown pathway with a single random time delay.^21-28^ This random time delay describes the time it takes between the signal activation and the production of response molecules through the unknown pathway, also known as signal transduction time (Figure 1A). The shape of the transduction-time distribution provides information about the underlying signaling pathway. For instance, the low mean and variance of the distribution indicates that the underlying pathway is fast and precise, respectively (Figure 1B). Furthermore, the number of modes in the distribution provides information about the structure of the underlying pathway.^25^ A unimodal transduction-time distribution appears when the response is regulated by a single-time-scale pathway (e.g., an irreversible chain and a reversible cascade) (Figure 1C-top). On the other hand, a transduction-time is multimodal when the response is regulated by multiple pathways with different transduction speeds, i.e., a multi-time-scale pathway (e.g., a cross-talk and feed-forward network) (Figure 1C-bottom). This indicates that inferring the shape of the transduction-time distribution can provide valuable information about the characteristics of the underlying signaling pathway.

**Figure 1.**
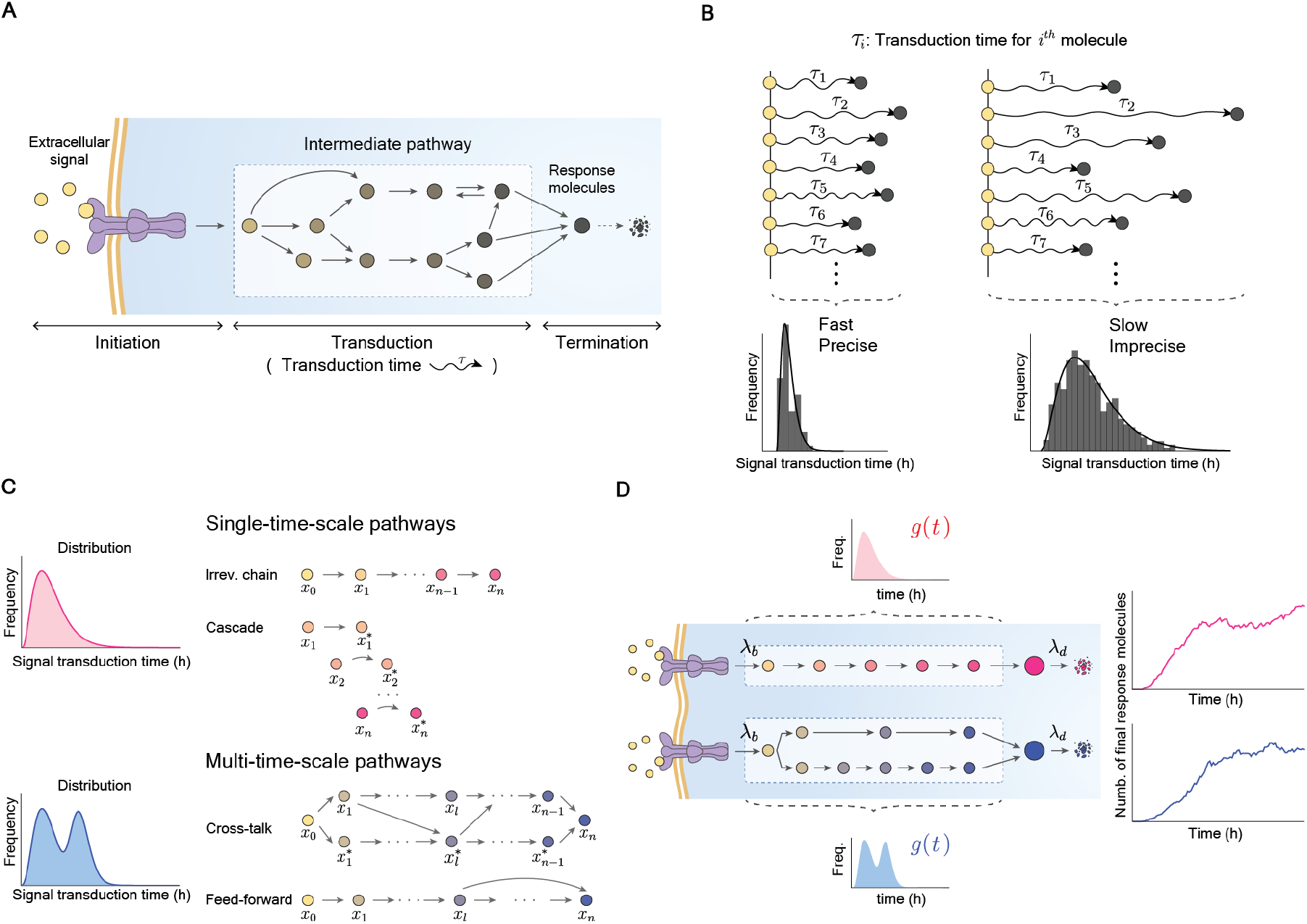
The distribution of signal transduction-time delay provides rich information about the underling lying signaling pathways. (A) The signal transduction time refers the time it takes between signal initiation and production of the final response molecules through intermediate pathways. (B) If the signal transduction is precise and fast (or imprecise and slow), then the distribution of the transduction-time (*τ*_*i*_) becomes narrow (or wide) and has a small (or large) mean. (C) Single- and multi-time-scale pathways have different number of modes of the transduction-time distribution. (D) The signaling pathways are modeled using a stochastic delayed birth-death process. The signal is activated at a rate of *λ*_*b*_ and transduced via signaling pathways. The response molecules are produced after the delay, whose distribution is *g*(*t*), and they decay at a rate of *λ*_*d*_. Although single- and multi-time-scale pathways have different transduction-time distributions (unimodal and multimodal), time traces of final product are indistinguishable.

Kim et al. developed a parametric Bayesian inference method that infers transduction-time distributions, revealing the rate-limiting steps in a signaling pathway.^21^ Through this approach, this study found that as the number of rate-limiting steps increases, so does cell-to-cell heterogeneity in response to antibiotic stress. However, this method only works if the underlying signaling pathway is a single-time-scale pathway, resulting in the transduction-time distribution following a Gamma distribution (Figure 1C-top). Similarly, other inference methods have limitations,^21-24^ as they are only applicable when sufficient information about the signaling pathway is available to specify the type of transduction-time distribution.

In this paper, we developed Density physics-informed neural network (Density-PINN), which infers the shape of transduction-time distributions in signaling pathways only from time traces of final stress response. Specifically, we modified PINN^29^ to incorporate physics-based knowledge of a signaling process with an arbitrary transduction-time distribution into the training of neural networks (NNs). We applied Density-PINN to single-cell gene expression time traces from 16 promoters in response to the antibiotic stresses, tetracycline (TET) and trimethoprim (TMP). This allowed us to uncover key features of unknown signaling pathways regulating these promoters, including the speed of signal initiation and transduction, transduction-time precision, and whether the promoter is regulated by single-or multi-time-scale pathways. Importantly, we found that promoters with longer signaling initiation and transduction time (i.e., longer response time) exhibit larger cell-to-cell heterogeneity in response intensity. However, this heterogeneity is greatly reduced when the response is regulated by multi-time-scale pathways (Figure 1C-bottom). This finding suggests that targeting pathways with shorter response times or involving multi-time-scale pathways can enhance the consistency of cellular responses and decrease unresponsive cells, which is critical for the development of anti-cancer drugs. Density-PINN provides an effective method to gain critical information about cell signaling pathways only from their response time traces.

## Results

### Cellular processes with hidden reactions can be described with a delayed model

Intracellular signaling pathways, activated by extracellular stimuli, can be described by a stochastic delayed birth-death process.^30-33^ In this model, the signal is activated at a rate of *λ*_*b*_, then it is transduced via signaling pathways and triggers the final response after a distributed time delay *g*(*t*), and the final response molecules decay at a rate of *λ*_*d*_ (Figure 1D). In this way, unobserved complex intermediate steps can be simply described with a delay distribution *g*(*t*). The transduction-time distribution *g*(*t*) is unimodal or multimodal depending on whether the underlying signaling pathways are single or multiple time scales (Figure 1C). The mean time trace of this stochastic process *y*(*t*) can be described by the following equation^21^ (see Note S1 for details):

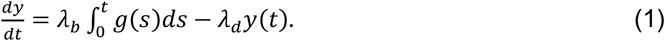

### Density-PINN: PINN-based estimation method for transduction-time distribution

As this formula provides a connection between the underlying transduction-time distribution *g*(*t*) and the final response *y*(*t*), it can be used to estimate the *g*(*t*) from the *y*(*t*). One promising approach for this purpose is to use PINNs, which are deep learning methods that integrate data and governing equations to estimate parameters. However, conventional PINNs can estimate parameter values rather than probability distribution.^34,35^ To address this problem, we propose Density-PINN that yields distribution 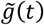 as the estimates of *g*(*t*) (see Note S7 for a step-by-step manual). Specifically, we used *M* Rayleigh distributions with different modes and widths as building blocks to construct the arbitrary distributions: 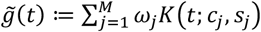, where *c*_*j*_ and *s*_*j*_ determine the mode and widths of Rayleigh distribution *K*(*t*; *c*_*j*_, *s*_*j*_) Thus, by estimating the parameters. (*ω*_*j*_, *c*_*j*_, *s*_*j*_) of 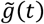, Density-PINN can estimate transduction-time distribution.

To perform efficient parameter estimation, we used evenly spaced shift parameters *c*_*j*_ within the time domain of the observed time series data *y*(*t*). Then, the remaining scale parameters *s*_*j*_ and weights *ω*_*j*_ were estimated through an artificial NN based on a Variational Autoencoder (VAE) (Figure 2A-VAE).^36,37^ Specifically, the artificial NN maps *y*(*t*) to a low-dimensional latent variable ***Z***, and generates distributions of *s*_*j*_ and *ω*_*j*_ from ***Z*** (see Methods for details). From the distributions of *s*_*j*_, *ω*_*j*_, and predetermined *c*_*j*_, a probability distribution of 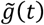 can be constructed (Figure 2A-Output).

**Figure 2.**
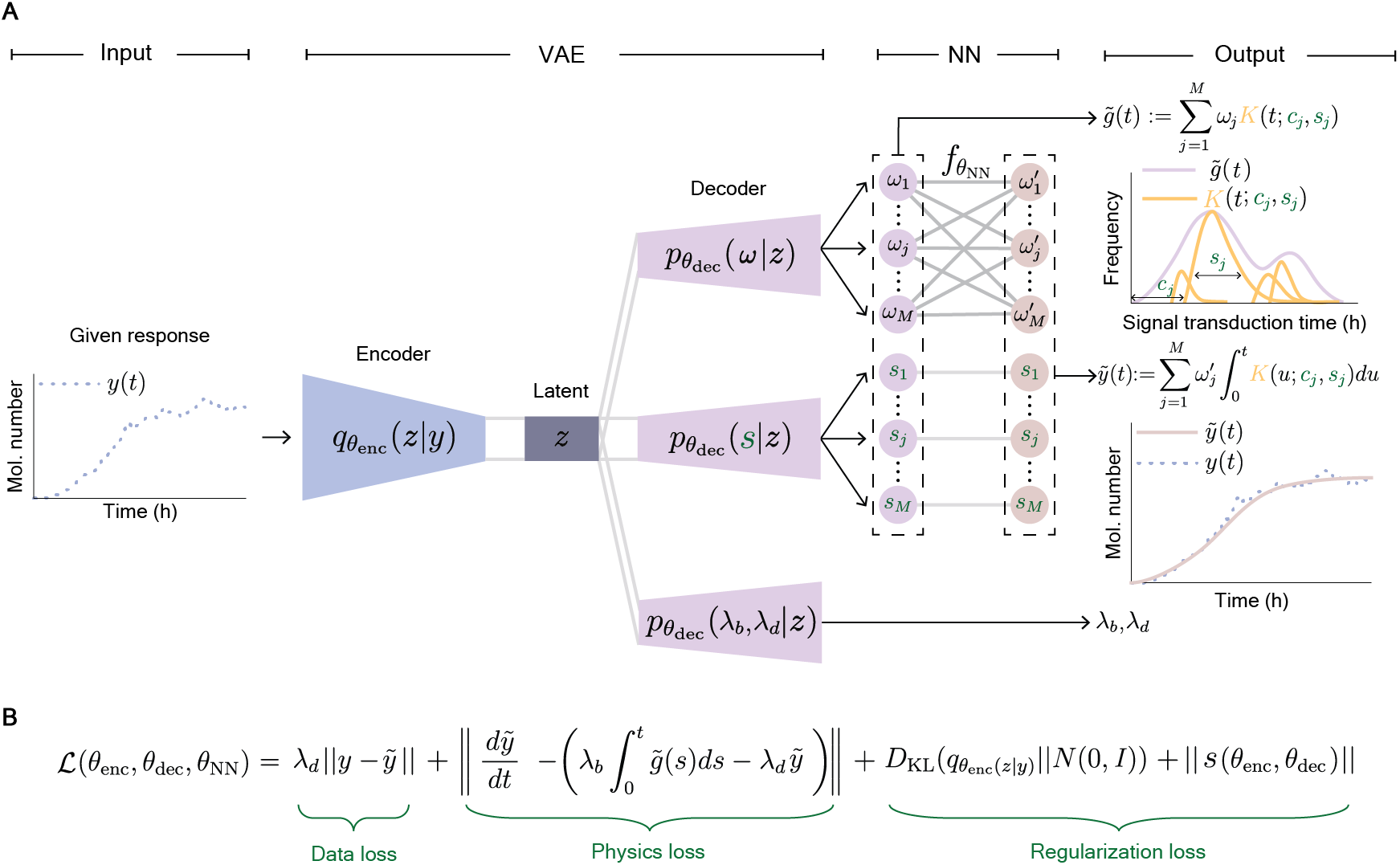
Schematic architecture of the framework of Density-PINN for inferring a delay distribution. (A) From the input response *y*(*t*) processed through VAE, a NN provides the estimated transduction-time distribution 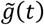, as well as the activation rate *λ*_*b*_, decay rate *λ*_*d*_, and the reconstructed response 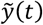. Specifically, in the VAE, the encoder 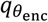, governed by the trainable parameters *θ*_enc_, maps *y*(*t*) to a latent variable ***Z*** in the latent space *Z*, and the decoder 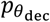, governed by the trainable parameters *θ*_*dec*_, maps the latent variable ***Z*** to a weight *ω*, scale parameter *s, λ*_*b*_, and *λ*_*d*_. The NN 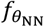, governed by trainable parameters *θ*_*NN*_, maps *ω* to the other weight *ω*^′^. The *ω*^′^, *ω* and *s* are used to compute 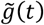 and 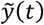 as 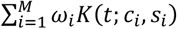 and 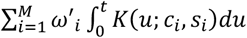, respectively, where *K*(*u*; *c*_*i*_, *s*_*i*_) is a shift Rayleigh densities with the shift parameter *c*_*i*_ and the scale parameter *s*_*i*_. (B) The total loss function is composed of the data loss, physics loss and regularization loss. The data loss quantifies the distance between *y*(*t*) and 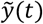. The physics loss quantifies how well 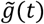 and 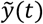 fit the governing equation (Equation 1). The regularization loss simultaneously prevents ***s*** from becoming too small or large and ensures effective representation of the data by the latent variables ***Z***.

To train 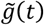 so that it approximates the true transduction-time distribution, *g*(*t*), we need to define a loss function. Because *g*(*t*) is unobservable, we cannot directly measure the difference between *g*(*t*) and 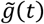. Instead, we indirectly quantified this difference by comparing the observed *y*(*t*) to the reconstructed 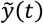 sobtained with 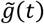 (Figure 2A-Output) (see Note S2 for details). To make sure that 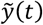 approximates *y*(*t*) and satisfies the governing equation (Equation 1), we used a data loss 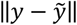 and physics loss 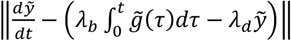 (Figure 2B). Since the physics loss contains *λ*_*b*_ and *λ*_*d*_, they were also estimated through a separate artificial NN (Figure 2A-Output). Furthermore, we included a regularization loss for scale parameters of *K*(*t*; *c*_*j*_, *s*_*j*_), ‖*s*‖ and a typical regulation loss of VAE, KL divergence *D*_KL_.(***Z*‖*N***(**O, *I***)). ‖*s*‖ prevent *s*_*j*_ from having too small or large values (see Note S3 for details), and *D*_KL_.***Z*‖*N***(**O, *I***)) ensures informative representation of data with latent variables.

For multiple input time traces {*y*_1_(*t*), *y*_2_(*t*), …, *y*_*N*_(*t*)}, the average of the total loss function (Figure 2B) of each time trace was minimized for the training of Density-PINN (Figure 3A-dashed box). Then, we estimated the distribution of *g(t), λ*_*b*_ and *λ*_*d*_ using the mean time trace 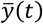 (Figure 3A).

**Figure 3.**
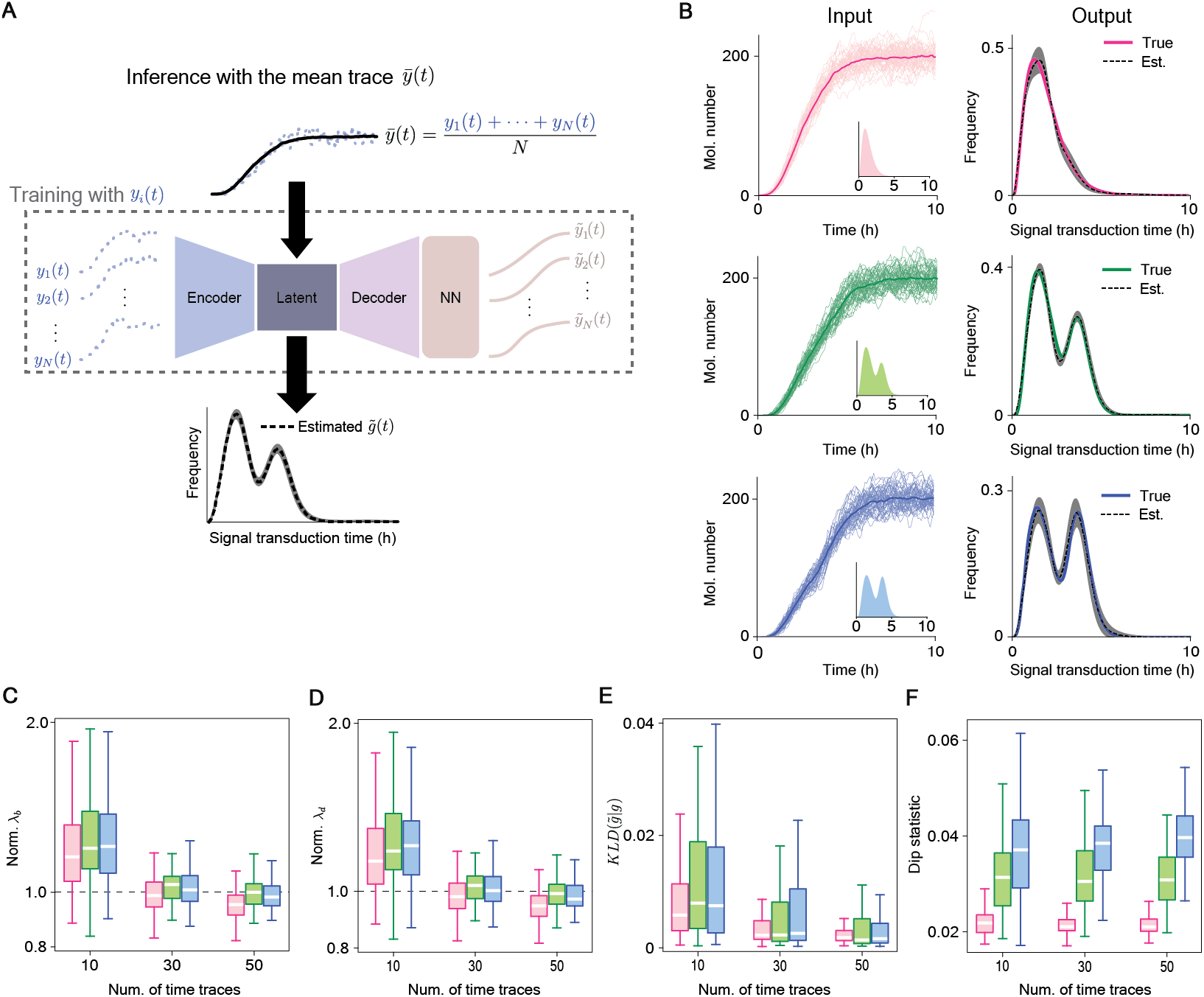
Density-PINN accurately estimates underlying transduction-time distributions with various shapes. (A) We trained Density-PINN with ***N*** individual response time traces so that the average of the total losses from ***N*** time traces is minimized. Using the trained model, a transduction-time distribution 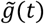 as well as *λ*_*b*_ and *λ*_*d*_ were inferred using the mean of the ***N*** time traces 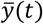, as an input. (B) Density-PINN accurately infers the underlying transduction-time distributions (Output) from simulated 50 time traces (Input) when the transduction-time distribution is unimodal (red), weakly bimodal (green), or strongly bimodal (blue) (Input; Inset). The shaded region represents the prediction intervals of the estimated transduction-time distribution (see Methods for details). Here, data was sampled every 0.5 hr. (C-F) As more time traces were used for the inference, the estimation becomes more accurate: estimations of the *λ*_*b*_ and *λ*_*d*_ become more accurate and precise (C, D), the KL-divergence between the underlying and reconstructed transduction-time distributions decreases (E), and the dip statistic, which increases as the bimodality increases, becomes more clearly distinguished among the unimodal (red), weakly bimodal (green), and strongly bimodal (blue) distributions (F).

### Density-PINN accurately estimates transduction-time distributions

We tested whether Density-PINN could estimate the three distinct transduction-time distributions: unimodal, weakly bimodal, and strongly bimodal distributions (Figure 3B-Input). We first generated 50 traces with each transduction-time distribution using a delayed stochastic simulation algorithm (see Note S5 for details).^38^ Then, we used these traces to estimate a transduction-time distribution 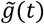, an activation rate *λ*_*b*_, and a decay rate *λ*_*d*_. Although the generated traces were nearly indistinguishable, our method successfully estimated the true transduction-time distributions for all three transduction-time distributions (Figure 3B-Output). In particular, the true transduction-time distributions are fully contained within the quantified prediction intervals (Figure 3B-Output, shaded region; see Methods for details).

We repeated this estimation 100 times by generating 100 different data sets, each containing 50 traces. Our method consistently provided accurate estimates of *λ*_*b*_ (Figure 3C) and *λ*_*d*_ (Figure 3D), the small distance between the true *g*(*t*) and the estimated 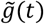 as quantified by KL-divergence (Figure 3E), and accurate estimates of bimodality as quantified by a dip statistic (Figure 3F) (see Note S6 for details). The accuracy and precision of the estimations improve as the number of time traces used for the estimations increases (Figures 3C-F).

When the decay mainly occurs via growth-induced dilution, the decay rate can be replaced with a dilution rate whose value can be estimated by measuring single-cell growth trajectories obtained with time-lapse microscopy. Thus, we tested our method when the decay rate *λ*_*d*_ was fixed to its true value. In this case, the estimations for the transduction-time distribution *g*(*t*) and the activation rate *λ*_*b*_ became more accurate and precise (Figure S2).

### Multi-time-scale pathways reduce the cell-to-cell heterogeneity in response

We applied our method to the previously measured single-cell time-lapse yellow fluorescent protein (YFP) expression data from 16 promoters in response to two antibiotic stresses, TET and TMP, in *Escherichia coli* (*E. coli*) populations (Figure 4A).^2^ The response time traces showed significant cell-to-cell heterogeneity. In particular, the final stress intensity is highly variable, which we quantified using a coefficient of variation (CV) calculated at the final observation point, referred to as the population CV of *y*. The population CV of *y* largely differed over the promoters (from 0.18 to 0.66; Figure 4A), even for the same antibiotic stress. However, it was unclear which properties of the signaling pathways affect the cell-to-cell heterogeneity in response due to the limited information available on the signaling pathways for each promoter.

**Figure 4.**
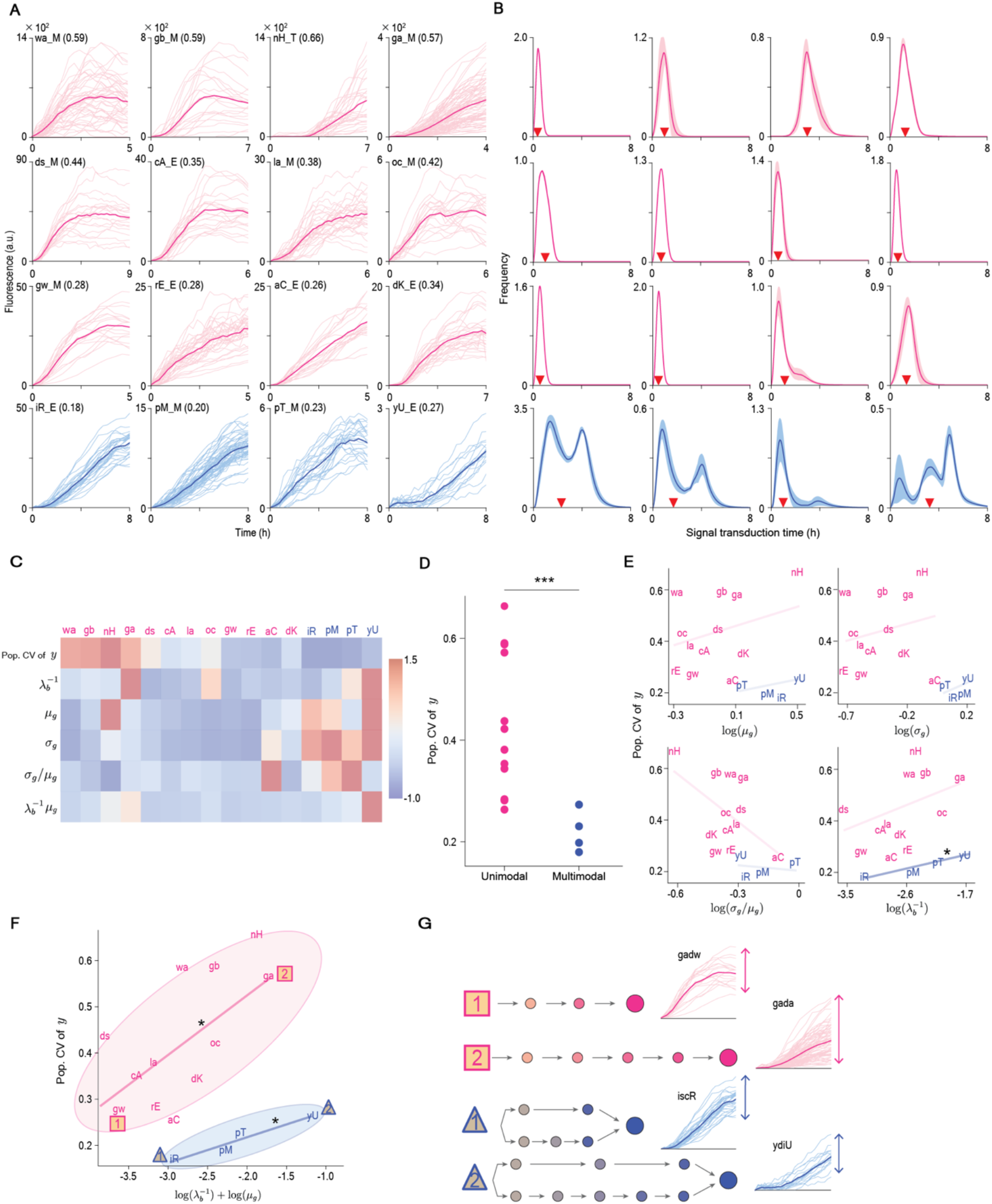
Multi-time-scale pathway leads to low cell-to-cell heterogeneity in response to antibiotic stresses. (A) Response time traces of 16 promoters to the antibiotic stresses, TET (E) and TMP (M), measured by time-lapse fluorescence microscopy.^2^ The thin and thick lines indicate single-cell time traces and the mean time traces, respectively. The numbers in parenthesis are the coefficient of variation (CV) of the responses at the final observation point. See Table S1 for the abbreviation of the promoters. (B) Estimated transduction-time distributions from the response time traces in Figure 4A. The estimated transduction-time distributions of twelve and the other four populations exhibit unimodality (red) and bimodality (blue), respectively. Red triangles represent the mean transduction times. (C) The characteristics of signaling pathways for each promoter quantified by Density-PINN: the initiation time 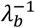, the mean ***µ***_*g*_, variance 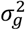, and the CV (=***σ***_*g*_/***µ***_*g*_) of transduction-time distribution. Additionally, we obtained the response time 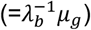 (see Method for details). Here, each quantity was standardized so that the mean and variance among the promoters are zero and one, respectively. (D) Population CVs of *y* are generally higher in the promoters with unimodal transduction-time distributions compared with those in the promoters with bimodal transduction-time distributions. Two-sided t-test was used for statistical test, *p* = 0.0003 (\*\*\**p* < 0.001). (E) None of the ***µ***_*g*_, ***σ***_*g*_, ***σ***_*g*_/***µ***_*g*_ and 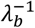 are significantly correlated with population CVs of *y*, except for 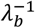 of multi-time-scale pathways (right bottom, blue). P-values were calculated by the Pearson correlation test: *p* = 0.164 (0.699), 0.605 (0.525), 0.055 (0.791), 0.063 (0.035), for ***µ***_*g*_, ***σ***_*g*_, ***σ***_*g*_/***µ***_*g*_, and 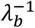 in single (multi)-time-scale-pathways, respectively. (F) On the other hand, the response time, 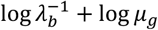 is significantly correlated with population CVs of *y* with *p* = 0.023 and 0.024 in single- and multi-time scale pathways. This correlation is smaller in multi-time-scale pathway. (G) Promoters with short response times (e.g., *gadw* and *iscR*) or involving multi-time-scale pathways (e.g., *iscR and ydiU*) show low level of heterogeneity.

To obtain information about the signaling pathways, we estimated the signal initiation time 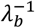, the time it takes to begin signal transduction in response to antibiotics, and the transduction-time distribution *g*(*t*) of each promoter, using Density-PINN. Throughout the estimation, the dilution rate, directly estimated from the experimentally measured cell growth rate ^2^, was used as the decay rate *λ*_*d*_ because dilution is the main driver of decay of YFP.^39^ Our inference results provided valuable information on the unknown signaling pathways regulating these promoters. Specifically, out of 16 promoters, 12 promoters exhibited unimodal transduction-time distributions (Figure 4B-red) while the other four promoters exhibited multimodal transduction-time distributions (Figure 4B-blue). This indicates that the 12 promoters are regulated by single-time-scale pathways (Figure 1C-top), while the other four promoters are regulated by multi-time-scale pathways (Figure 1C-bottom). Furthermore, the mean ***µ***_*g*_, the standard deviation ***σ***_*g*_, and the CV (=***σ***_*g*_/***µ***_*g*_) of each transduction-time distribution *g*(*t*) across the promoters reveals the speed and precision of the signal transduction time (Figure 4C).

We next investigated the relationship between these quantified characteristics of signaling pathways and the cell-to-cell heterogeneity (i.e., population CV of *y*). Interestingly, the cell-to-cell heterogeneities in response to promoters regulated by single-time-scale pathways were higher compared to those regulated by multi-time-scale pathways (Figure 4D). However, none of the other characteristics, such as the means, standard deviations, and CV of transduction-time distributions, were significantly correlated with the cell-to-cell heterogeneity in response (*p* > 0.05, Figure 4E) except for the signal initiation time 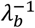 in multi-time scale pathways (Figure 4E-right bottom). In particular, it was unexpected that a large variation in transduction-time distribution (i.e., a highly variable timing of response) did not lead to a large cell-to-cell heterogeneity in final response intensity.

Unlike the signal initiation time 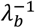 and the mean of transduction time ***µ***_*g*_, interestingly, their sum, i.e., the response time, is significantly correlated with the cell-to-cell heterogeneity in both single- and multi-time-scale pathways (Figure 4F). The positive relationship was stronger in the single-time-scale pathway group compared with the multi-time-scale pathway group. In particular, when the response time was short (e.g., for gadw and iscR promoters), the cell-to-cell heterogeneity in response to antibiotic stress was small (Figures 4F and 4G-rectangle and triangle 1). When the response time was long (e.g., for gada and ydiU promoters), the cell-to-cell heterogeneity was large if the response is regulated by a single-time-scale pathway (Figures 4F and 4G-rectangle 2) while the heterogeneity was still relatively small if the response is regulated by a multi-time-scale pathway (Figures 4F and 4G-triangle 2). Interestingly, this result is consistent with previous findings that multi-time-scale pathways reduce cell-to-cell heterogeneity in cell volume recovery,^18^ which was concluded from intensive experimental work using mutant yeast cells.

## Discussion

We developed Density-PINN that accurately infers parameter values and transduction-time distribution of a stochastic process with time delay (Figures 2 and 3). We applied Density-PINN to single-cell time-lapse fluorescent protein expression data in response to two antibiotic stresses, TET and TMP (Figures 4A-4C). This uncovered key properties of the signaling pathways leading to the cell-to-cell heterogeneity in stress response: an increase in heterogeneity with longer response time (Figure 4F) and a decrease in heterogeneity when triggered by multi-time-scale pathways (Figure 4G). These results highlight the importance of response time and pathways, which can be inferred with our method, in identifying effective target molecules for drug development (Figure 5). Our findings also enable a systematic understanding of the heterogeneity of treatment effects, which is a major challenge for precision medicine.^40^

**Figure 5.**
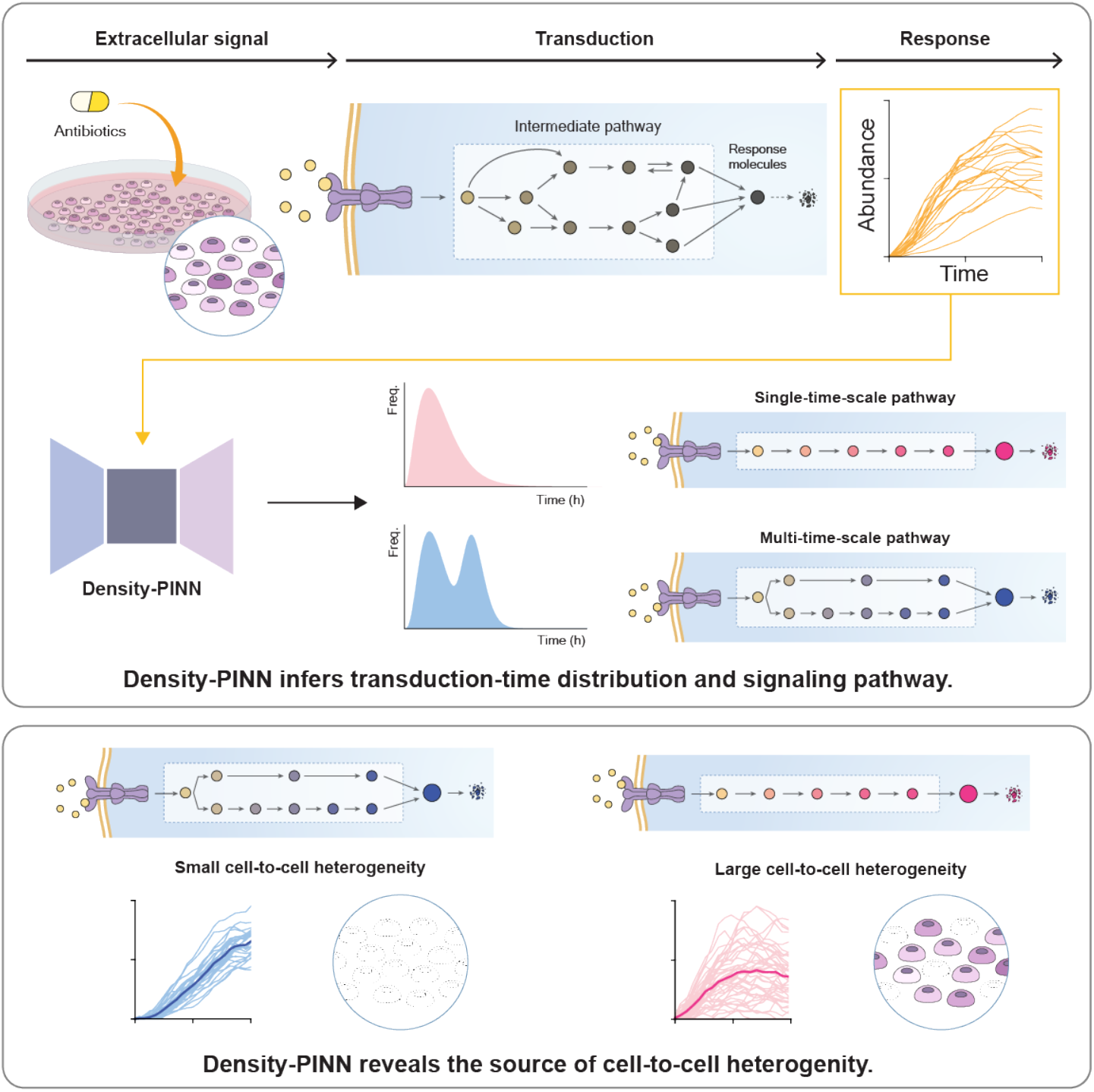
Summary Figure. Density-PINN reveals the source of cell-to-cell heterogeneity by inferring transduction-time distributions. (Top) When *E. coli* cells are exposed to antibiotic stress, responding proteins are produced via signaling pathways. From the time trace of the accumulated responding proteins, Density-PINN infers the distribution of signaling transduction time, whose shape informs the number of pathways with different time scales. (Bottom) This revealed that the presence of signaling pathways with different time scales greatly reduce the cell-to-cell heterogeneity in response to antibiotics.

We illustrated the application of Density-PINN with a focus on cases where the final response increases and then saturates, which can be described by Equation 1. For more complex dynamics, such as adaptation or oscillation profiles, the mean formula of the stochastic delayed birth-death process with feedback regulation, derived in our previous work,^21^ can be utilized instead of Equation 1. Then, by simply adjusting the physics loss term based on the mean formula,^41,42^ Density-PINN can be applied to analyze signaling pathways with a wider range of dynamics.

Recently, PINNs have emerged as a powerful tool for inferring parameter values of differential equations because it incorporates prior physical knowledge within NNs while accurately fitting data. In this study, we proposed Density-PINN that estimates a probability distribution as well as parameter values. Density-PINN produces outputs that naturally conform to the properties of probability distributions by using the kernel density estimation. In this way, additional loss terms to impose constraints on the probability distributions, causing additional computational costs, was not needed unlike previous approaches.^34,35^ To improve computational efficiency further, we chose the Rayleigh distribution as the kernel density since it satisfies the vanishing condition at time *t* ≤ 0 and allows direct calculation of definite integration of the estimated distribution in the physics loss without numerical integration.

PINNs have demonstrated their versatility in solving a wide variety of problems across different fields of science and engineering.^41^ PINN have been originally used to solve deterministic differential equations^29,34,43^ and extended its application to optimize engineering designs,^44^ solve inverse problems^45,46^, and stochastic differential equation.^47-49^ In this study, we pioneer the use of PINN in the analysis of non-Markovian systems with distributed time delays, which can be applied to a variety of fields. For example, Density-PINN can be used to infer the distribution of time delays due to the latent period of COVID-19^50^ (Figure S4). Additionally, Density-PINN can be used to calculate a lower bound of entropy production rate of a system by inferring a residence time distribution in each state of a non-Markovian model.^51^ It is particularly important as the entropy production rate quantifies the extent to which a cell is consuming energy in order to resist environmental changes.^52^

## Methods

### Formulation of Density-PINN architecture

*Y* = {***y***_1_, ***y***_2_, …, ***y***_*N*_} is the set of ***N*** time traces whose mean and standard deviation are ***µ*** (*t*) and ***σ***_*Y*_(*t*), respectively. The standardized *Y* by replacing *y*_*i*_(*t*) with (*y*_*i*_(*t*) – ***µ***_*Y*_(*t*))/ ***σ***_*Y*_(*t*) was used as input of the encoder of the VAE. The encoder then mapped each standardized ***y***_*i*_ to two parameters ***µ***_*i*_ and 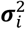, used to parametrize the lower-dimensional latent variable ***Z***_*i*_ = ***µ***_*i*_ + ***σ***_*i*_ × ***ε*** where ***ε***∼***N***(**0, *I***):

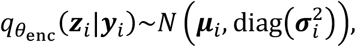

where 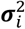 is the elementwise square of ***σ***_*i*_. The dimension of ***Z***_*i*_ (*k*) is 4 and is much smaller than the dimension of ***y***_*i*_ (*d*) used in this study. Thus, the encoder consists of one input layer of size *d*, two hidden layers of size 16, and two output layers of size *k* = 4 for ***µ***_*i*_ and log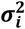. The goal of the encoder is to train the parameter *θ*_enc_ such that 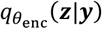 is as close as possible to the true posterior distribution.

The decoder of the 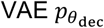 with the trainable parameter *θ*_*dec*_ consists of three fully connected NNs. The first two networks have the same architecture with two hidden layers of size 16, and the last network has one hidden layer of size 16. The three NNs transform the latent variable ***Z***_*i*_ to the weights ***ω***_*i*_, the scale parameter ***s***_*i*_, and the activation and decay rates (*λ*_*b,i*_, *λ*_*d,i*_) to obtain the estimated transduction-time distribution 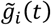 with predefined shift parameters {*c*_1_, …, *c*_*M*_} as follows:

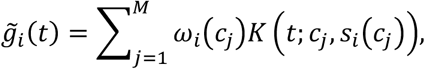

where *K* (*t*; *c*_*j*_, *s*_*i*_.*c*_*j*_)) is a shifted Rayleigh density with the scale parameters *s*_*i*_(*c*_*j*_) and the shift parameters *c*_*i*_ weighted by *ω*_*i*_(*c*_*j*_) (Figure 2A-bottom). That is, 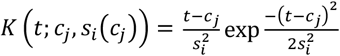 for *t* ≥ *c*_*j*_ and *K* (*t*; *c*_*j*_, *s*_*i*_.*c*_*j*_)) = 0 otherwise. We chose the Rayleigh distribution rather than negative binomial^36^ or Gaussian distribution^37^ used in previous studies as it has positive support. Moreover, this choice allowed the definite integration of 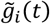 in the physics loss, which reduces the computational cost and numerical error of numerical integration. See Table S2 for the hyperparameters in Density-PINN.

The VAE whose output is 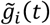 cannot be directly trained because the true transduction-time distribution *g*_*i*_(*t*) is unobservable. Thus, we used 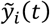 reconstructed from 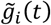 and compared it with the true ***y***_*i*_ (Figure S1). To do this, we employed a NN, 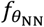, with trainable parameters *θ*_*NN*_ that consists of one input, hidden, and output layer whose size of same as the length of ***ω***_*i*_. Thus, the NN mapped weights ***ω***_*i*_ to new weights 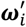 for reconstructing 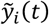 as follows:

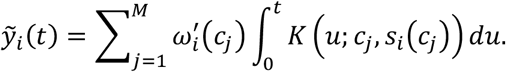

### Training of Density-PINN

For the training, we used the Adam optimizer^53^, and to prevent overfitting issues, we applied an early stopping criterion^54^ (see Note S4 for details). Density-PINN was trained by minimizing the total loss function:

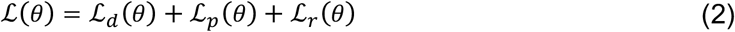

where *θ* = (*θ*_enc_, *θ*_*dec*_, *θ*_*NN*_), ℒ_*d*_ is a data loss, ℒ_*p*_ is a physics loss, and ℒ_*r*_ is a regularization loss. The data loss function is defined as:

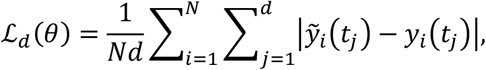

where {*t*_1_, …, *t*_*d*_} is the set of observed time points of *y*_*i*_(*t*) The physics loss is evaluated the set of collocation points, 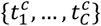, evenly spaced in the time domain [0, *T*]:

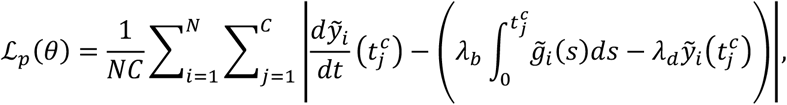

where the derivative of 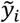 with respect to time *t* is computed using automatic differentiation.^55^ The regularization loss is computed as follows:

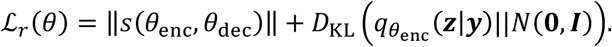

The first term is given by

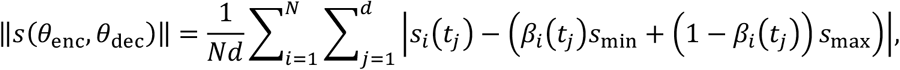

where 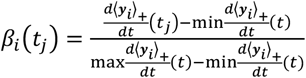 and *s*_min_ and *s*_min_ are the lower and upper bounds of the scale parameters, respectively. This term is used to smoothen 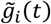 (see Note S3 for details). Here 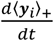 denotes the nonnegative part of numerical derivative of ***y***_*i*_ after applying moving average with the window size of *L* = 7. The second term 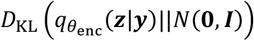 is a typical regulation loss of VAE used to ensure informative representation of data with latent variables (Figure S3).

### Inference using Density-PINN

After training Density-PINN, we obtained the mean and standard deviation, (***µ, σ***) for the latent variable ***Z***, by passing the average of time traces 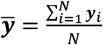 through the encoder. Using (***µ, σ***), we generated 1000 latent samples .***Z***^(1)^, ***Z***^(2)^, …, ***Z***^(1000)^) from a normal distribution ***N***(***µ***, diag(***σ***^2^)). Each sample, ***Z***^(*l*)^ was then passed through the decoder, resulting in 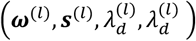. With these parameters, we constructed an estimated transduction-time distribution 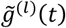 as follows:

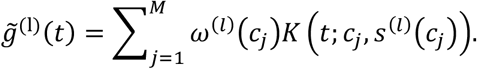

We calculated the sample mean 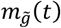 and standard deviation 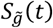 of 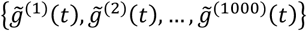. We then obtain the upper boundary 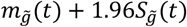 and the lower boundary 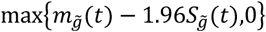, i.e., 95% prediction interval. The shaded regions in Figures 3B and 4B indicate the areas between the boundaries.

### **Interpretation of the response time**, 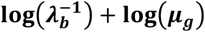

The inverse of activation rate 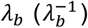 represents the signal initiation time per each molecule as the unit of *λ*_*b*_ is the number of molecules per time. However, the unit of observed response molecules is often not the number of molecules but the unit of fluorescence. In this case, the estimated 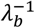 has the following relationship with the true initiation time 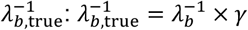 where the *γ* is a conversion rate from the number of molecules to the unit of fluorescence. Thus, to convert 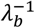 to 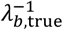, we need *γ*, which can be obtained by measuring the binomial error in partition of total YFP signal during cell division.^56^ However, *γ* is unknown in the experimental data we used in Figure 4. Therefore, we used log.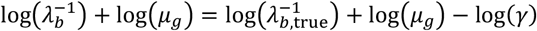 in Figure 4F. Since this is the sum of signal initiation time and transduction times in log scale, shifted by log(*γ*) (i.e., shifted response time in log scale), the observed positive associations in Figure 4F are preserved for the true response time without shift.

## ACKNOWLEDGEMENTS

We thank Life Science Editors for editorial support and Sunghwan Bae (Bstar Artwork) for scientific illustration. H.H. is supported by the National Research Foundation of Korea (NRF) NRF-2019-Fostering Core Leaders of the Future Basic Science Program/Global Ph.D. Fellowship Program 2019H1A2A1075303. H.J.H. is supported by the National Research Foundation of Korea (NRF) grant funded by the Korea government (MSIT) (No. RS-2022-00165268). W.C. is supported by the Charles Phelps Taft Research Center at the University of Cincinnati (No. M80941). J.K.K. is supported by the Institute for Basic Science IBS-R029-C3 and Samsung Science and Technology Foundation SSTF-BA1902-01.

## AUTHOR CONTRIBUTIONS

H.J., H.H., H.J.H., and J.K.K. designed the study. H.J. developed an algorithm. H.J., H.H., W.C., and J.K.K. analyzed results. H.J., H.H., and J.K.K. wrote the manuscript. All authors revised the manuscript.

## DECLARATION OF INTERESTS

The authors declare no competing financial interests

## Notes

### Competing Interest Statement

The authors have declared no competing interest.

